## Supplementary information for "Darwinian Evolution of Self-Replicating DNA in a Synthetic Protocell"

for

### Content

Pages 2-4 **Supplementary Notes**

Page 2 **Supplementary Note 1:** Replicator engineering

Page 2 **Supplementary Note 2:** Reducing the set of proteins for self-replication in liposomes

Page 3 **Supplementary Note 3:** Error-prone IVTTR with an exonuclease-deficient DNAP variant

Page 3 **Supplementary Note 4:** Mock selection of an active self-replicator

Page 4 **Supplementary Note 5:** Characterization of purified DNAP protein variants

Pages 5-6 **Supplementary Tables**

Page 5 **Table S1.** Codon bias analysis on at least 5% synonymous mutations from all evolutionary campaigns.

Page 5 **Table S2**. Primer pairs used for PCR and qPCR

Page 6 **Table S3.** Transitions of the MS/MS measurements for the proteolytic peptides of DNAP

Page 6 **Table S4.** Mutations found by Sanger sequencing of twelve single clones isolated from round 11 of the Int-WT(1) evolution campaign.

Pages 7-21 **Supplementary Figures**

Pages 7-21 **Figs S1 – S16**

Page 22 **Supplementary References**

### Supplementary Notes

#### Supplementary Note 1: Replicator engineering

Sanger sequencing of the gel-purified 1.4 kb DNA fragment revealed that this band corresponded to a shortened *ori-p3* fragment that lost the DNAP gene-expression cassette (T7 promoter-gene-T7 terminator), but still had the origins and the entire TP transcriptional unit intact. Thus, it could potentially act as a potent parasite. We hypothesized that the deletion of the DNAP cassette was a result of a recombination event between two identical 83-bp DNA sequences on the *ori-p2p3* template that spanned the T7 promoter, the following T7 g10 leader, and a ribosome binding site upstream the two genes. To diminish the recombination frequency during IVTTR, we modified the repeated T7 g10 leader sequence upstream of the TP gene on the template **(Fig. 1c)**. We hypothesized that the strong stem-loop structure that the original leader sequence forms on the transcribed mRNA, as predicted by RNAfold web server[1], could be substituted with an artificial sequence with a similar mRNA secondary structure for avoiding recombination without affecting gene expression of TP **(Fig. 1c).** We therefore constructed a modified *ori-p2p3* template wherein 33 bp of the leader sequence upstream of the *p3* gene was replaced by an alternative sequence that was predicted to form a similar mRNA stem loop despite no primary sequence similarity to the original leader sequence. This resulted in a modified *ori-p2p3* template (we called *mod-ori-p2p3*) with a similar TP expression **(Fig. S1a)** and self-replication ability as the original *ori-p2p3* **(Fig. 1b, Fig. S1b)**, although it seems to produce additional smaller DNA products of sizes roughly 1.1 and 2.5 kb. These additional DNA fragments are not visible after in-liposome IVTTR followed by PCR recovery using primers that bind to the extreme ends of *mod-orip2p3* (**Fig. 1b**). The original 1.4 kb DNA fragment from *ori-p2p3* is likewise barely detectable after in-liposome IVTTR and PCR recovery in *mod-orip2p3* (**Fig. 1b**). The absence of 1.1 and 2.5 kb DNA fragments after PCR amplification suggests that these are likely unfinished DNA products produced during bulk IVTTR amplification of *mod-orip2p3*, rather than molecular parasites with two origins of replication capable of displaying significant competition to replicators. Together, these data corroborate the recombination hypothesis and the idea that the secondary RNA structure of the g10 leader sequence, and not its primary sequence, is somehow crucial for gene expression. Lastly, they suggest that *mod-ori-p2p3* is a better starting template for our in vitro evolutionary experiments.

#### Supplementary Note 2: Reducing the set of proteins for self-replication in liposomes

We sought to optimize the compartmentalised IVTTR reaction. We have previously shown that both SSB and DSB are required in high amounts for efficient self-amplification of *ori-p2p3* in bulk IVTTR reactions[2]. However, we also observed that in an orthogonal DNA amplification setting (i.e., where the amplified DNA encodes a protein other than DNAP and TP), a reduced amount of DSB improved the yield of expressed protein from the replicating template[3]. We therefore investigated if reducing DSB concentration could improve self-amplification in liposomes. Surprisingly, we found that *mod-ori-p2p3* self-amplifies efficiently in liposomes even in complete absence of DSB **(Fig. S2)**. However, consistent with bulk IVTTR that strongly relies on the presence of DSB, template DNA outside of liposomes did not amplify without DSB **(Fig. S6)**. Therefore, we chose to omit DSB in our follow up evolution protocol.

**Supplementary Note 3: Error-prone IVTTR with an exonuclease-deficient DNAP variant**

We explored the position F62 **(Fig. 1d)** (originally reported as F65, due to an erroneous delimitation of the translation initiation site. Our delimitation corresponds to the NCBI Reference Sequence: NC_011048.1). The F62 residue is located in the N-terminal exonuclease domain of DNAP **(Fig. 1d)**, and is involved in stabilising at the active site the ssDNA that is formed when the template/primer DNA melts during proofreading[4]. The F62Y mutation was reported to reduce exonucleolytic activity of DNAP and increase the frequency of nucleotide misincorporation, while only mildly affecting the DNAP's TP-deoxynucleotidylation, TP-DNA initiation, and TP-DNA amplification[4]. To test TP-primed DNA amplification activity by the F62Y variant of DNAP, we first constructed a plasmid containing the circularised *mod-ori-p2(F62Y)p3* (hence it cannot self-replicate) and performed orthogonal DNA amplification of an origin-flanked unrelated gene, *ori-pssA* (the *pssA* gene encodes for *E. coli* phosphatidylserine (PS) synthase), in a bulk IVTTR reaction. Absolute quantification of the *pssA* gene by quantitative polymerase chain reaction (qPCR) before and after 16 hours of reaction revealed a yield of amplified DNA comparable to that performed by wild-type (WT) DNAP **(Fig. 1e)**. Secondly, we compared the in-liposome self-replication activity of *mod-ori-p2p3* and *mod-ori-p2(F62Y)p3*, and found that both versions reached similar amplification folds in our system **(Fig. S1c)**. We therefore decided to also use this polymerase variant, along with the wild-type DNAP, in some of our evolution experiments**.**

**Supplementary Note 4: Mock selection of an active self-replicator**

We performed an enrichment experiment starting from a mock library consisting of *mod-ori-p2p3* and a 50-fold molar excess of an unrelated gene (*plsB*, coding for *E. coli* glycerol-3-phosphate acyltransferase) of similar length and flanked with Φ29 replication origins, *ori-plsB* (**Fig. S3a**). To ensure a stringent genotype-to-phenotype link, we encapsulated an estimated 0.2 DNA molecules per liposome on average ($\lambda$ = 0.2) - considering an average vesicle diameter of 4 µm and a Poissonian partitioning[3], this corresponds to a total DNA concentration of 10 pM in solution. To prohibit IVTTR outside of liposomes, the DNA outside of liposomes was digested with externally added DNase I. After 16 hours of in-liposome IVTTR at 30 °C, we measured the concentration of *p2* and *plsB* genes before and after incubation by absolute qPCR quantitation, which revealed that *mod-ori-p2p3* fraction in the

DNA population increased approximately 10-fold after a single round of in-liposome IVTTR **(Fig. S3b)**. These results imply that our developed strategy of in-liposome IVTTR of a self-replicator can support an in vitro evolution campaign.

**Supplementary Note 5: Characterization of purified DNAP protein variants**

To disentangle protein property from DNA template effects, we assayed the replication activity of the purified S79G, A80T, and the double mutant DNAP variants. Under different replication conditions, with or without coupled transcription, the DNA template harbouring the S79G mutation was not amplified better than the WT DNA template by purified WT or DNAP variants **(Fig. S15a-c)**. This result indicates that S79G alone did not improve the template replicability either. Curiously, we also observed that both purified S79G and A80T DNAP variants fail to replicate DNA in a Φ29 replication buffer **(Fig. S15d,e)**, both in protein-primed and DNA-primed settings **(Fig. S16)**. The mutated residues are located in the 3'-5' exonuclease domain of DNAP. Since this domain also contains residues involved in stabilizing either the primer terminus at the active centre[5,6] or the template strand[5–7], we suspected a defect in the interaction of the mutant proteins with DNA in certain conditions. To test this idea, we measured the affinity of the mutant proteins to the template/primer complex by electrophoretic mobility shift assay and observed reduced DNA binding **(Fig. S16)**. This result suggests that the replication deficiency the mutants display in specific conditions, such as in the Φ29 replication buffer, and/or when replication is initiated by a DNA-oligomer primed mechanism, may be explained by the impaired stabilisation of the 5' end of the template strand. Nevertheless, it is not clear if such destabilisation of the DNAP-DNA complex could improve the protein-primed DNA replication in a higher ionic-strength and crowded environment, such as the conditions we employed in our self-replication system.

### Supplementary Tables

**Table S1.** Mutations found by Sanger sequencing of twelve single clones isolated from round 11 of the Int-WT(1) evolution campaign.

| Clone | Gene/Sequence | Mutation | Original codon | New codon |
|---|---|---|---|---|
| 1 | *p2* (DNAP) | S79G | AGC | GGC |
| | | V247 silent | GTG | GTA |
| | | E311K | GAA | AAA |
| | | V562A | GTT | GCT |
| | *p3* (TP) | R70 silent | CGT | GCG |
| | | L263P | CTG | CCG |
| | T7 terminator | | CTA | CTG |
| | | | AAC | ACC |
| 2 | *p2* (DNAP) | S79G | AGC | GGC |
| | | V247 silent | GTG | GTA |
| | | G294 silent | GGC | GGT |
| | *p3* (TP) | Y175 silent | TAC | TAT |
| | | L263P | CTG | CCG |
| | T7 terminator | | CTA | CTG |
| 3 | *p2* (DNAP) | S79G | AGC | GGC |
| | | P261 silent | CCG | CCA |
| | *p3* (TP) | K188 silent | AAA | AAG |
| | T7 terminator | | CTA | CTG |
| 4 | *p2* (DNAP) | I67 silent | ATT | ATC |
| | | A80T | GCC | ACC |
| | | N85S | AAC | AGC |
| | | V247 silent | GTG | GTA |
| | | Q443 silent | CAA | CAG |
| | *p3* (TP) | K79N | AAA | AAT |
| | | K246 silent | AAA | AAG |
| | T7 terminator | | CCC | CCA |
| | | | GGG | GGA |
| 5 | *p2* (DNAP) | S79G | AGC | GGC |
| | | G141 silent | GGC | GGT |
| | | V247 silent | GTG | GTA |

| Clone | Gene/Sequence | Mutation | Original codon | New codon |
|---|---|---|---|---|
| 6 | *p2* (DNAP) | S79G | AGC | GGC |
| | | M243V | ATG | GTG |
| | | K256 silent | AAA | AAG |
| 7 | *p2* (DNAP) | S79G | AGC | GGC |
| | | C103 silent | TGC | TGT |
| | | V247 silent | GTG | GTA |
| | | W274 stop codon | TGG | TAG |
| 8 | *p2* (DNAP) | S79G | AGC | GGC |
| | | A171 silent | GCA | GCG |
| | | V247 silent | GTG | GTA |
| | | V342I | GTT | ATT |
| | *p3* (TP) | S196 silent | TCG | TCA |
| 9 | *p2* (DNAP) | S79G | AGC | GGC |
| | | V247 silent | GTG | GTA |
| | | G414 silent | GGC | GGT |
| | *p3* (TP) | K188 silent | AAA | AAG |
| 10 | *p2* (DNAP) | S79G | AGC | GGC |
| | | V247 silent | GTG | GTA |
| | *p3* (TP) | R6H | CGC | CAC |
| 11 | *p2* (DNAP) | D54N | GAC | AAC |
| | | L83 silent | CTG | TTG |
| | | I199V | ATC | GTC |
| | | V247 silent | GTG | GTA |
| | *p3* (TP) | A137T | GCA | ACA |
| | | K188 silent | AAA | AAG |
| 12 | *p2* (DNAP) | C19 silent | TGC | TGT |
| | | S79G | AGC | GGC |
| | | A171T | GCA | ACA |
| | | V247 silent | GTG | GTA |
| | *p3* (TP) | L263P | CTG | CCG |
| | T7 terminator | | CTA | CTG |

**Table S2.** Codon bias analysis on at least 5% synonymous mutations from all evolutionary campaigns. Numeric fractions for codon usage estimation on each codon were taken from GenScript Codon Usage Frequency Table (https://www.genscript.com/tools/codon-frequency-table)

| Evolution campaign (at least 5% frequency) | AA position | Codon in parental *mod-orip-2p3* | Codon in original Φ29 genome | DNA mutation | Comments |
|---|---|---|---|---|---|
| Int-WT(1) | I67 | ATT (0.49) | ATC (0.39) | ATC (0.39) | Recovered Φ29 original DNA sequence |
| | V247 | GTG (0.35) | GTT (0.28) | GTA (0.17) | |
| | N248 | AAT (0.49) | AAT (0.49) | AAC (0.51) | Changed from original Φ29 sequence |
| Int-WT(2) | L477 | CTG (0.35) | TTG (0.13) | CTT (0.12) | |
| Int-Mut | I67 | ATT (0.49) | ATC (0.39) | ATC (0.39) | Recovered Φ29 original DNA sequence |
| | K121 | AAA (0.74) | AAG (0.26) | AAG (0.26) | Followed by string of A's |
| | K475 | AAA (0.74) | AAG (0.26) | AAG (0.26) | Followed by string of A's |
| Con-WT | E158 | GAA (0.68) | GAA (0.68) | GAG (0.32) | Changed from original Φ29 sequence |

**Table S3**. Primer pairs used for PCR and qPCR

| Primer pair set | Sequence (5' → 3') | Purpose |
|---|---|---|
| 1058 ChD<br>1060 ChD | GTCGACTGCTAATACGACTCACTATAGGGCCCTCTGGAGACACCAGAGGG<br>CGGGCTGCGTGCCATTAGTATATCTCCTTCTTAAAGTTAAACAAATAAACATGT | Cloning G340 plasmid containing *mod-ori-p2p3*. PCR fragment containing modified T7 leader sequence |
| 1049 ChD<br>1056 ChD | GCAAGGCGATTAAGTTGGGTAACG<br>TGTCTCCAGAGGGCCCTATAGTGAGTCGTATTAGCAGTCGAC | Cloning G340 plasmid containing *mod-ori-p2p3*. PCR fragment containing *p2* gene |
| 1057 ChD<br>1052 ChD | ACATGTTTATTTGTTTAACTTTAAGAAGGAGATATACTAATGGCACGCAGCCCG<br>TGTGTGGAATTGTGAGCGGATAAC | Cloning G340 plasmid containing *mod-ori-p2p3*. PCR fragment containing *p3* gene |
| 948 ChD<br>1132 ChD | TGTAAAACGACGGCCAGT<br>GTTGATAATGAACGCACCATCGTATTTCAGATTGTGGAAGTAC | Cloning G371 plasmid containing *mod-ori-p2p3* encoding for Φ29 DNAP(F62Y). Fragment 1 for Gibson assembly |
| 1131 ChD<br>1137 ChD | GTACTTCCACAATCTGAAATACGATGGTGCGTTCATTATCAAC<br>GGCGGTCATGCGATCCAG | Cloning G371 plasmid containing *mod-ori-p2p3* encoding for Φ29 DNAP(F62Y). Fragment 2 for Gibson assembly |
| 1324 ChD<br>1325 ChD | ATGGGGCGCCGATGGTCTGCCGAACACC<br>TCGGCGCCCCATTTAAAGCCATTACGTTCCAG | Cloning G559 plasmid containing *mod-ori-p2(S79G)p3*. |
| 1326 ChD<br>1327 ChD | CTGAAGAAACTGCCGTTTCCGGTGAAG<br>GTTTCTTCAGGCTGTCATAGATCACGGTATG | Cloning G560 plasmid containing *mod-ori-p2(K121K)p3*. |
| 1328 ChD<br>1329 ChD | GAAGAAACTGGGTTATTGGGCACACGAATC<br>GTTTCTTCGGGTCAACGATATCTTTAATCACATCC | Cloning G561 plasmid containing *mod-ori-p2(K475K)p3*. |
| 1330 ChD<br>1331 ChD | ATCAAGAGCGTCGAAGGCTCATTTAACTCGTTC<br>ACGCTCTTGATAAAATTCAGTTGCAGCTGAATC | Cloning G562 plasmid containing *mod-ori-p2p3(K188K)* |
| 1344 ChD<br>1345 ChD | ATGGGGCACCGATGGTCTGCCGAACACC<br>ATCGGTGCCCCATTTAAAGCCATTACGTTCC | Cloning G569 plasmid containing *mod-ori-p2(S79G&A80T)p3* |
| 1346 ChD<br>1347 ChD | TGGAGCACCGATGGTCTGCCGAACACC<br>CATCGGTGCTCCATTTAAAGCCATTACGTTCC | Cloning G570 plasmid containing *mod-ori-p2(A80T)p3* |
| 491 ChD<br>492 ChD | P-AAAGTAAGCCCCCACCCTCACATG<br>P-AAAGTAGGGTACAGCGACAACATACAC | PCR to produce *ori-p2p3*, *mod-ori-p2p3*, and all *mod-ori-p2p3* reversed engineered versions. |
| 976 ChD<br>977 ChD | GGATGAAGACTACCCGCTGC<br>ACAGGTCTGCGATTTCACCG | qPCR amplicon targeting *p2* gene |
| 365 ChD<br>952 ChD<br>953 ChD<br>982 ChD<br>1068 ChD | CAGTCACGACGTTGTAAAACGAC<br>GGCGATATTGACTATCACAAAGAACG<br>GATGTTACCGGTAAAGTCCCGTACC<br>GACAAAGCCGAATACGCCCG<br>CCATGATTACGCCAAGCTTGCATGC | Sanger sequencing of twelve single clones isolated from round 11 of Int-WT(1) |

**Table S4.** Transitions of the MS/MS measurements for the proteolytic peptides of DNAP

| Protein | Proteolytic peptide | Precursor ion *(m/z)* | Product ion *(m/z)* | Ion name |
|---|---|---|---|---|
| DNAP | ENGALGFR | 432,2221 | 492,2929 | y4 |
| DNAP | ENGALGFR | 432,2221 | 563,3300 | y5 |
| DNAP | ENGALGFR | 432,2221 | 620,3515 | y6 |
| DNAP | ENGALGFR | 432,2221 | 734,3944 | y7 |
| TP | IAEIER | 365,7083 | 314,1710 | b3 |
| TP | IAEIER | 365,7083 | 304,1615 | y2 |
| TP | IAEIER | 365,7083 | 417,2456 | y3 |
| TP | IAEIER | 365,7083 | 546,2882 | y4 |
| TP | IAEIER | 365,7083 | 617,3253 | y5 |

### Supplementary Figures

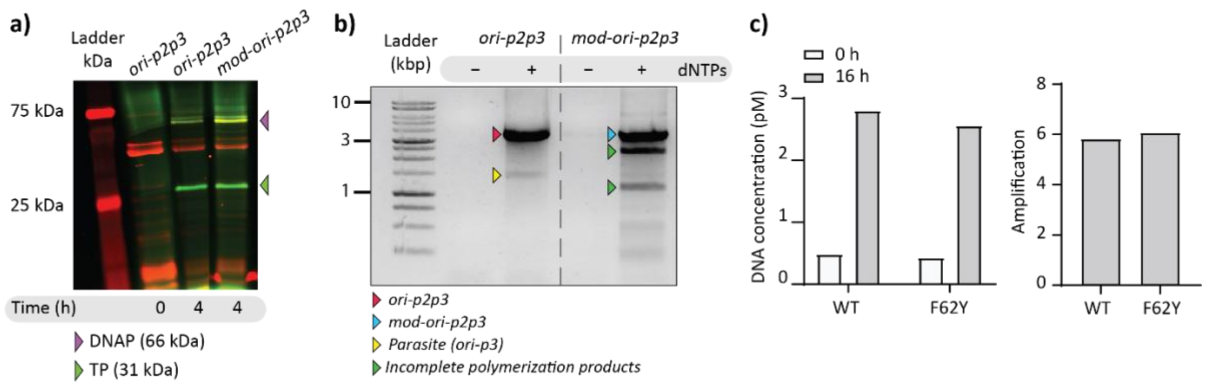


**Fig. S1: Characterization of an engineered DNA self-replicator. a)** Protein production from bulk IVTT reactions with *ori-p2p3* and *mod-ori-p2p3* DNA template. Each reaction solution was supplemented with Green-Lys reagent for co-translational protein labelling. Synthesized DNAP and TP were visualised by SDS-PAGE and fluorescence gel imaging. **b)** Agarose gel electrophoresis of recovered DNA from bulk IVTTR with *ori-p2p3* or *mod-ori-p2p3* DNA template. **c)** Assay of self-replication of *mod-ori-p2p3* expressing WT or F62Y DNAP in in vesiculo IVTTR reaction. Concentration of *p2* gene was quantified by qPCR (left), showing similar amplification levels (right) with both polymerases.

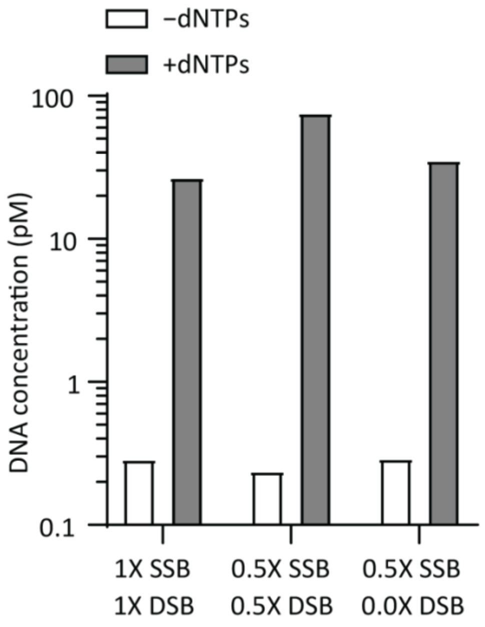


**Fig. S2: Protein DSB is dispensable in in-liposome IVTTR.** Absolute DNA quantification from in-liposome IVTTR reactions with different amounts of DSB and SSB. Omitting DSB does not hamper self-replication. Amplification by qPCR targeted the *p2* gene of the *mod-ori-p2p3* DNA construct. Initial concentration of *mod-ori-p2p3* DNA was 10 pM.

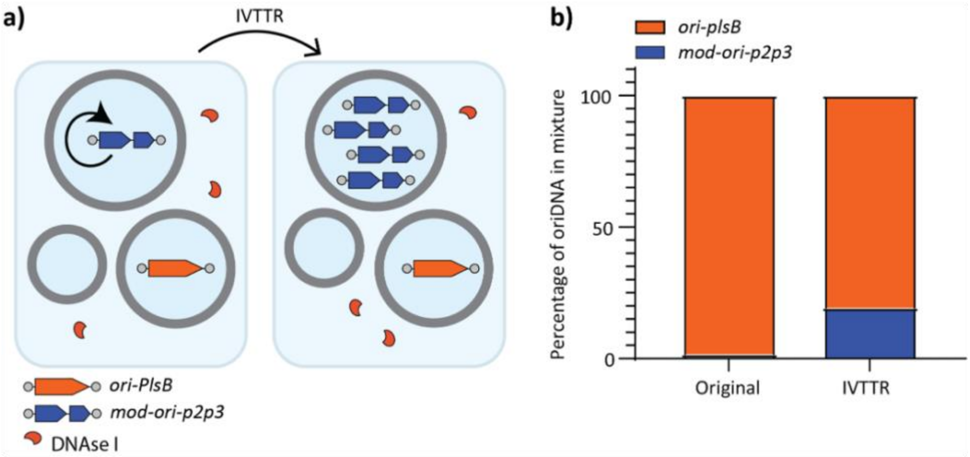


**Fig. S3: Clonal DNA self-amplification in liposomes**. **a)** Scheme of *mod-ori-p2p3* enrichment from a mixture with non-self-amplifying DNA (*ori-plsB*) inside gene-expressing liposomes. Total DNA concentration at the start of IVTTR was 10 pM, with a 50-fold excess of *ori-plsB* over *mod-ori-p2p3*. **b)** Calculated percentages of each DNA species in the original DNA mixture and after IVTTR. Enrichment of *mod-ori-p2p3* indicates that self-amplification is clonal.

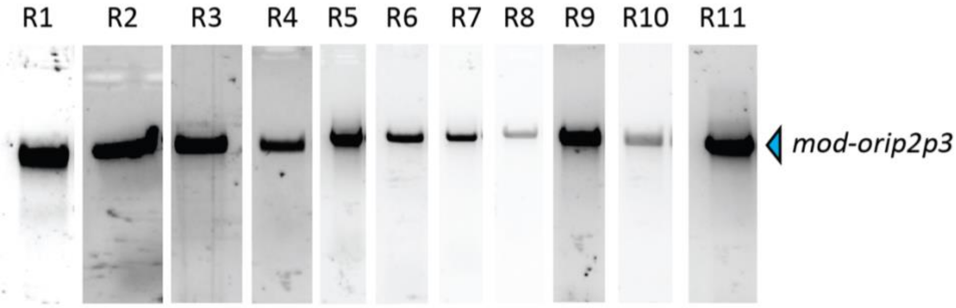


**Fig. S4: Agarose gel of Int-WT(2) samples after PCR recovery.** Only the full-length *mod-ori-p2p3* template can be seen at each evolution round. Therefore, DNA was not gel-extracted to begin a next IVTTR reaction.

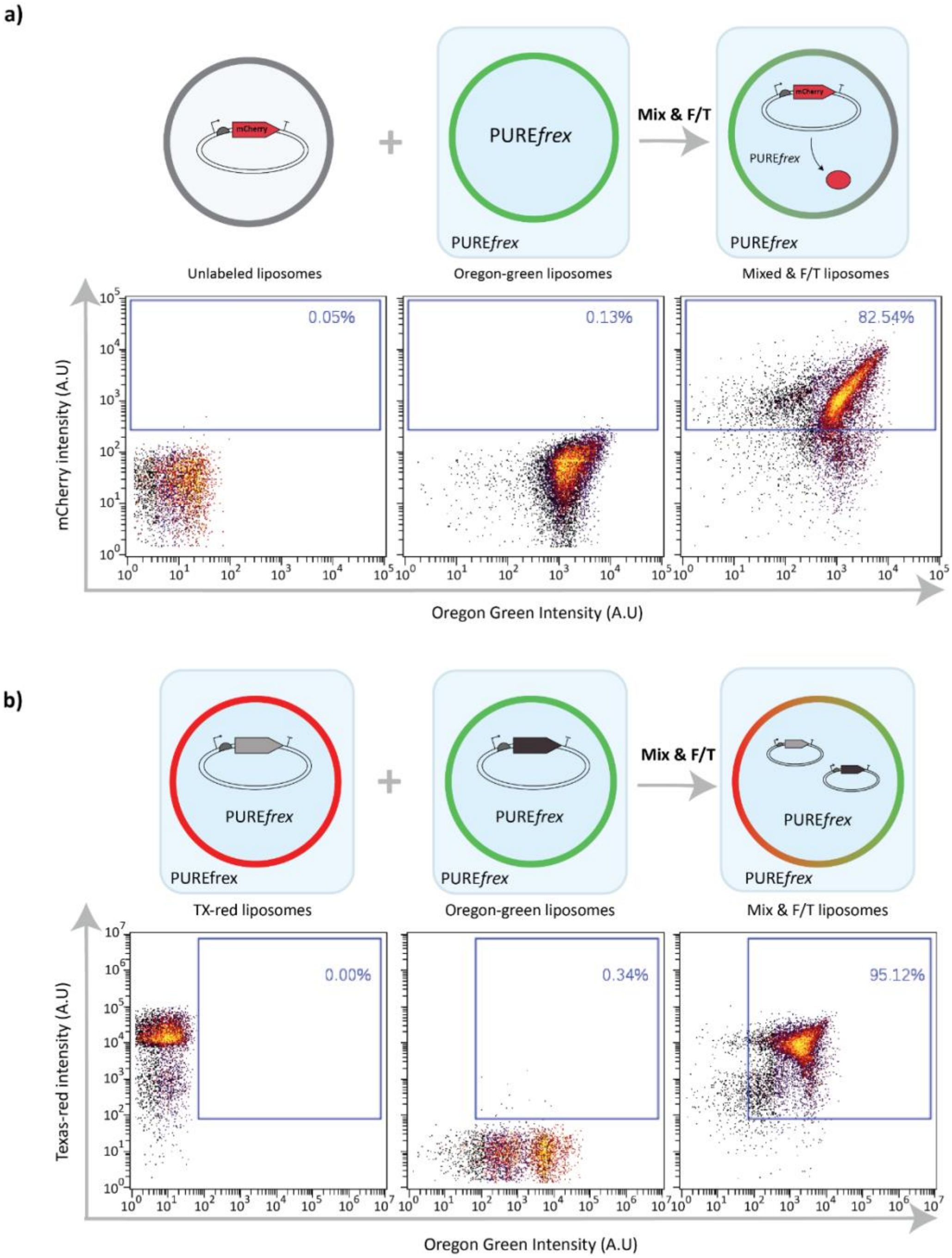


**Fig. S5: Liposome content mixing and membrane fusion are promoted by F/T. a)** Flow cytometry scatter plots from unlabelled liposomes encapsulating the mCherry gene (left), Oregon-green labelled liposomes with encapsulated PURE system (middle), and mixed samples exposed to F/T (right). The three liposome samples were incubated for 3 hours at 37 °C. The shift of mCherry fluorescence intensity toward higher values shows content mixing of the two liposome populations, which resulted in the production of mCherry. The appended percentage values indicate the percentages of mCherry-positive liposomes. **b)** Flow cytometry scatter plots from PURE-containing liposomes stained either with either Texas Red (left) or Oregon-green (middle) membrane dye. After liposome mixing and F/T (right), a new population of liposomes exhibiting both fluorophores (percentage values are appended) appeared as a result of lipid mixing.

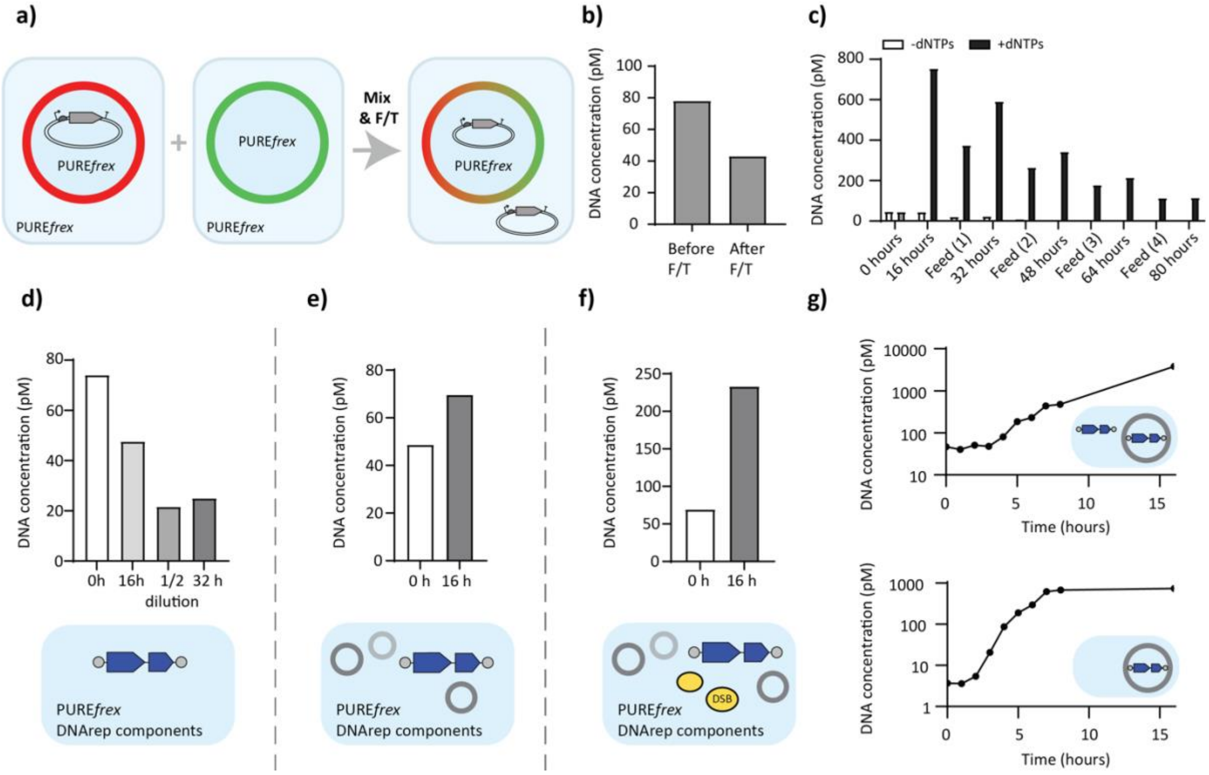


**Fig. S6: Design of the semi-continuous evolution experiment. a)** DNA leakage caused by F/T was assayed by measuring the DNA concentration after mixing and F/T of two liposome populations, one containing PURE and DNA, while the other contained only PURE. DNA outside of liposomes was digested by DNase I and the quantity of DNA was assessed by qPCR before and after F/T. **b)** Absolute DNA quantification reveals that ~50% DNA is lost (released outside liposomes) after F/T. **c)** Repeated in-liposome IVTTR reactions with *mod-ori-p2p3* DNA (semi-continuous evolution method) and no DSB protein (also no DNase I). A two-fold dilution with feeding vesicles was applied in between rounds. Omitting DSB causes a gradual decrease in DNA concentration. **d)** DNA *mod-ori-p2p3* cannot self-replicate in bulk IVTTR reactions without DSB protein. Samples were taken at different time points, with ½ dilution with fresh PURE after 16 hours incubation, and another 16 hours of incubation. **e)** DNA *mod-ori-p2p3* cannot efficiently self-replicate outside preformed liposomes without DSB. **f)** Addition of DSB enables self-replication outside liposomes. **g)** Kinetics of IVTTR reactions containing DSB with *mod-ori-p2p3* DNA present both inside and outside of liposomes (top), or only inside (bottom). IVTTR is more effective when compartmentalized in liposomes than outside.

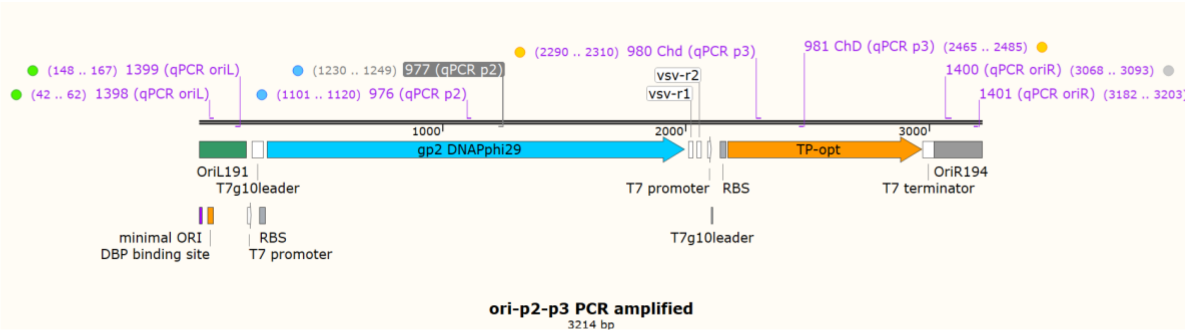


**Fig. S7: Schematic illustration of *mod-ori-p2p3* self-replicator displaying the regions that were targeted by qPCR.** Primer pairs were designed to target *oriL*, *p2*, *p3*, and *oriR* regions. The corresponding sequences (start and end) are indicated in brackets, as well as the primer names.

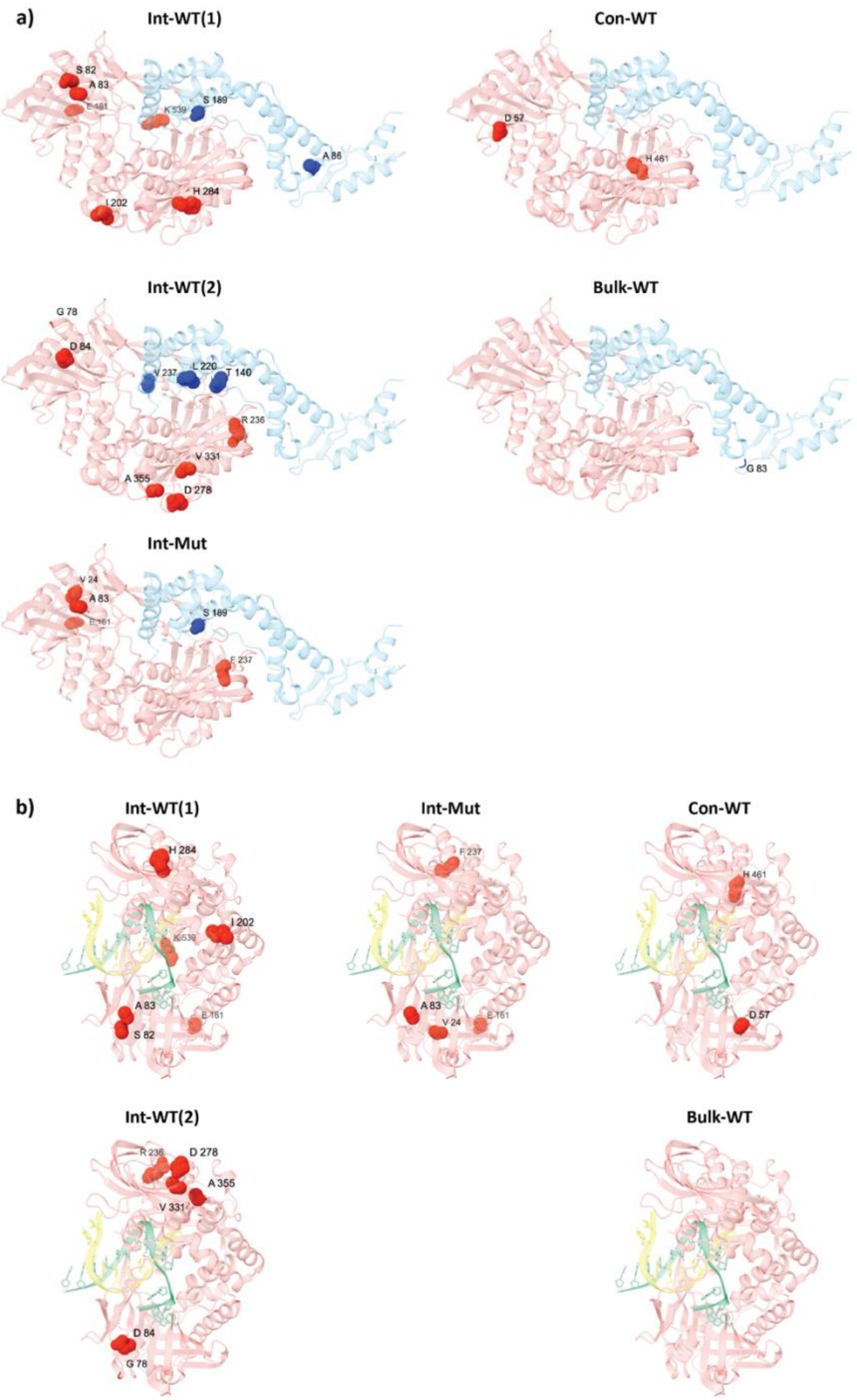


**Fig. S8: Visualisation of DNAP and TP protein residues on which mutations were detected at frequencies of at least 5%. a)** Residues in DNAP-TP protein complex (PDB 2EX3) are highlighted. **b)** Residues in DNAP primer-template complex (PDB 2PZS) are highlighted. DNAP is coloured in pink, TP in sky blue, DNA strand in green, primer in yellow, DNAP amino acid residues in red, and TP amino acid residues in blue. Molecular graphics and analyses were performed with UCSF ChimeraX[8]. Evolutionary campaigns are Int-WT(1), Int-WT(2), Int-Mut, Con-WT, and Bulk-WT.

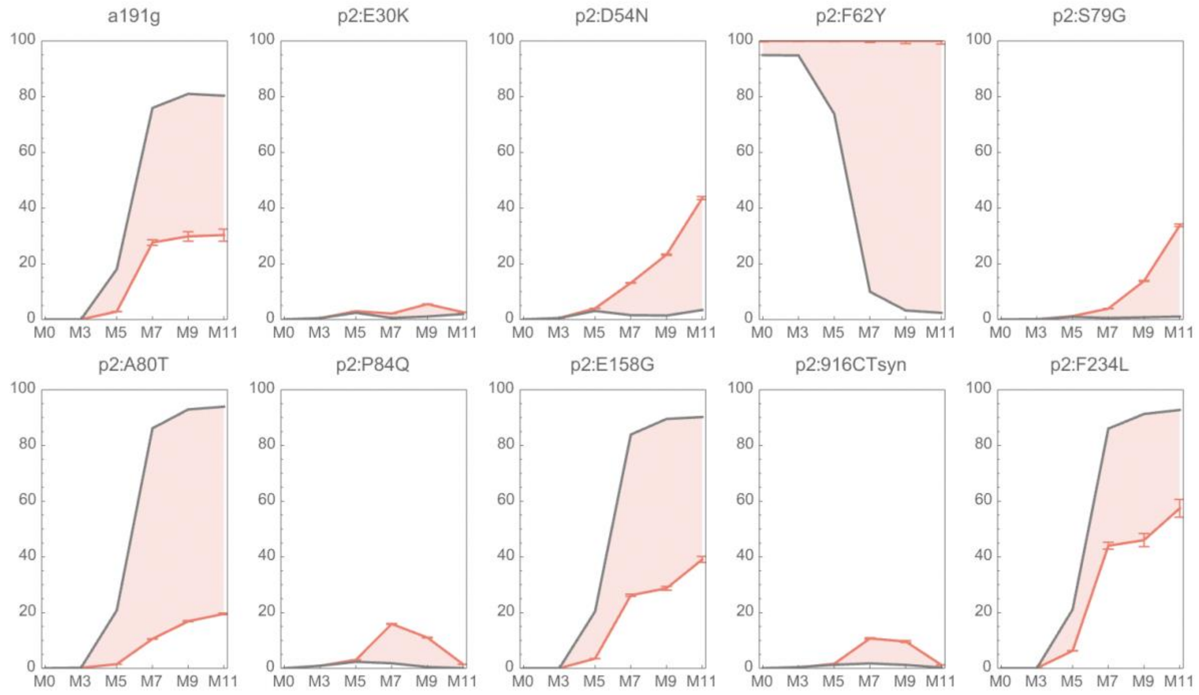


**Fig. S9: Dynamics of the mutation frequency in DNA variants containing the mutation F62Y from the Int-Mut evolution campaign.** The plots show that fraction of paired-end reads containing the mutation indicated above each plot, among all reads in grey and among reads that include P2F62Y in red. The mutation p2:F62Y was gradually removed from the library (grey line in the p2:F62Y panel), while substitutions p2:D54N and p2:S79G were strongly (and independently) enriching in the F62Y population (red lines in the corresponding panels), outcompeting an initial increase in p2:P84Q. Other mutations that increase (but are less frequent than in the general population) are probably the result of recombination events. Mutations associated to the invading DNA (without F62Y) include a191g, p2:A80T, p2:E158G, and p2:F234L, and they behave as a group. Rounds of evolution are indicated as M0, M3,…M11.

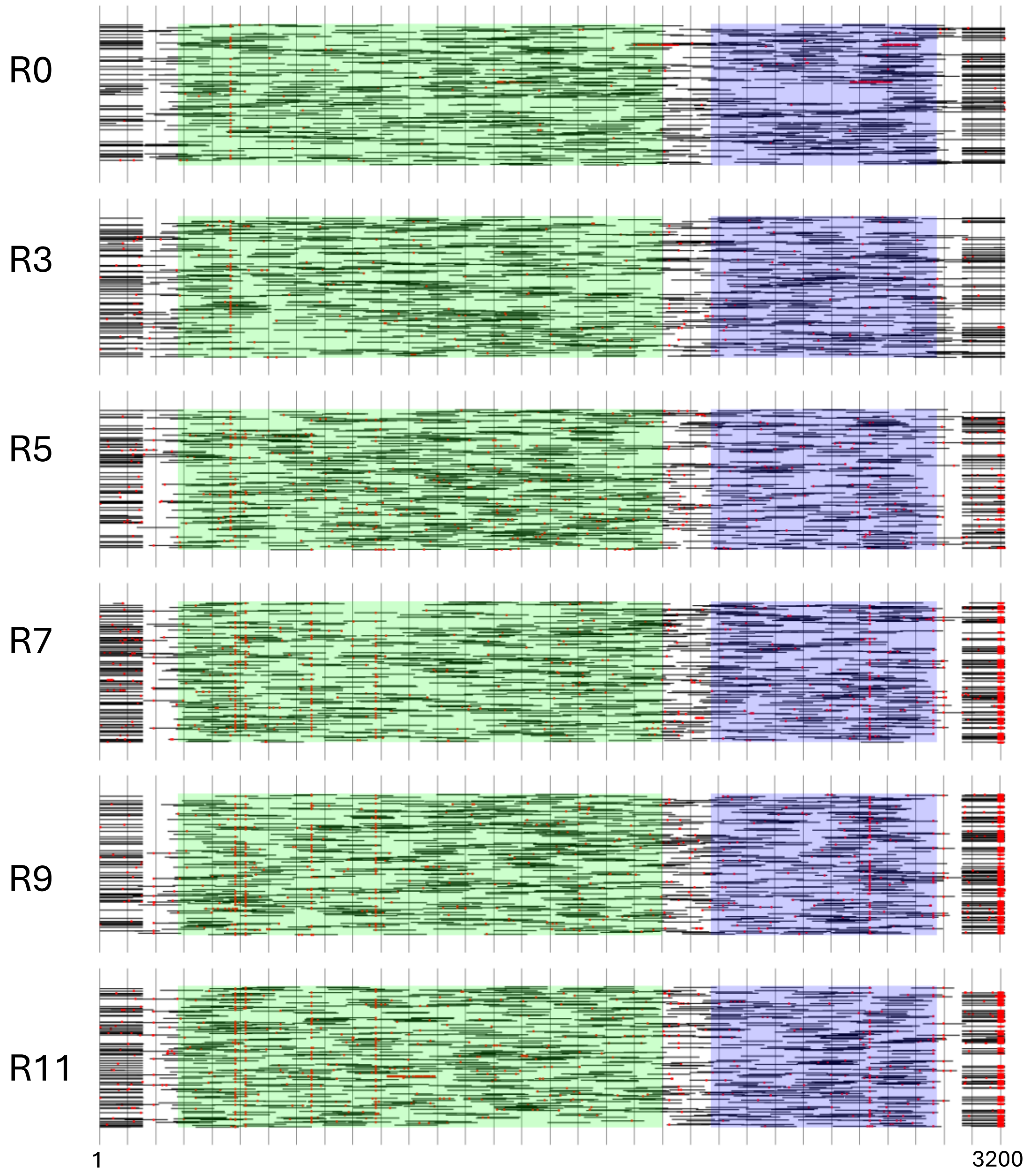


**Fig. S10: Paired-end read alignment of *mod-ori-p2(F62Y)p3* evolution on parental DNA shows reshaping of oriR during Int-Mut evolution campaign.** The IVTTR round is indicated to the left. Only 500 paired-end reads are shown in each round. Single-nucleotide mutations are shown as red dots. Green, *p2* CDS: Purple, *p3* CDS. Reshaping of oriR started at round 3, resulting from the swapping of oriL primer site to oriR.

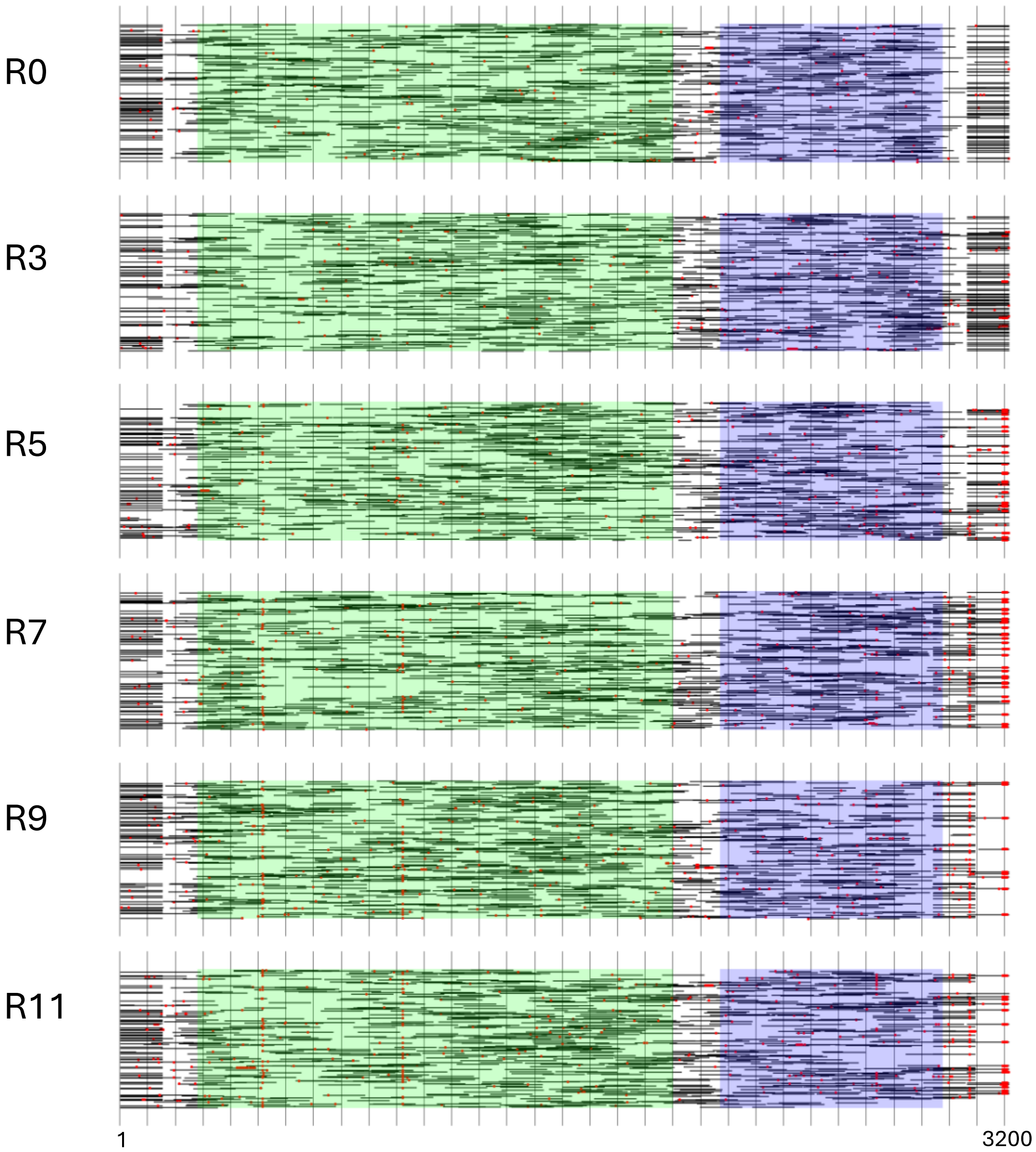


**Fig. S11: Paired-end read alignment of *mod-ori-p2-p3* on parental DNA shows reshaping of oriR during Int-WT(1) evolution campaign.** The IVTTR round is indicated to the left. Only 500 paired-end reads are shown in each round. Single-nucleotide mutations are shown as red dots. Green, *p2* CDS: Purple, *p3* CDS. Global rearrangements of oriR result from the swapping of oriL primer site to oriR (starts at round 3), followed by a large deletion at oriR (starts at round 5).

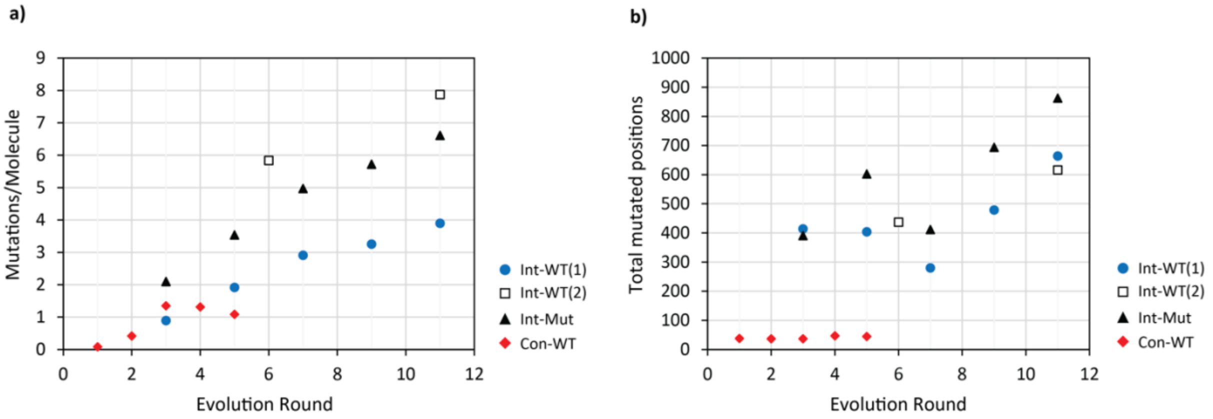


**Fig. S12:** NGS sequencing reveals accumulation of mutations per round of evolution. The number of mutations per molecule **a)** and total mutated positions **b)** are reported for all in-liposome evolutionary campaigns. Intermittent evolution: Int-WT(1), Int-WT(2), and Int-Mut. Continuous evolution: Con-WT.

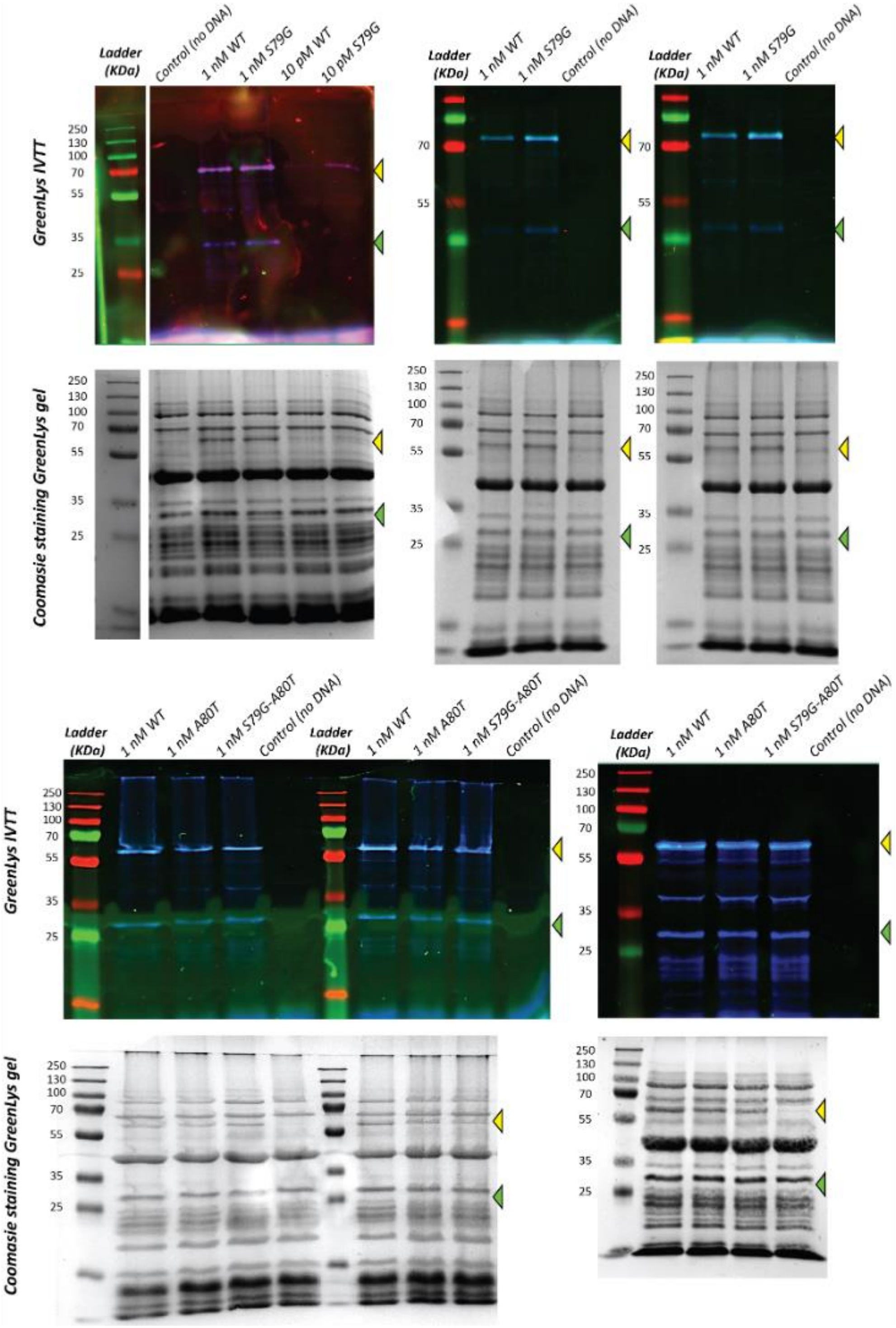


**Fig. S13: Expression of parental and reverse engineered *mod-ori-p2p3* DNA results in similar levels of synthesized proteins.** Bulk IVTT reactions were carried out with supplemented Green-Lys protein labelling reagent. The parental *mod-ori-p2p3* or reversed engineered variants (DNAP(S79G) and DNAP(A80T)) were used as DNA templates, as indicated. SDS-PAGE gels from six repeats are shown. For each repeat, the Green-Lys fluorescence image (upper gel) and Coomassie protein staining image (bottom gel) are displayed. The yellow and green arrowheads indicate the DNAP and TP bands, respectively.

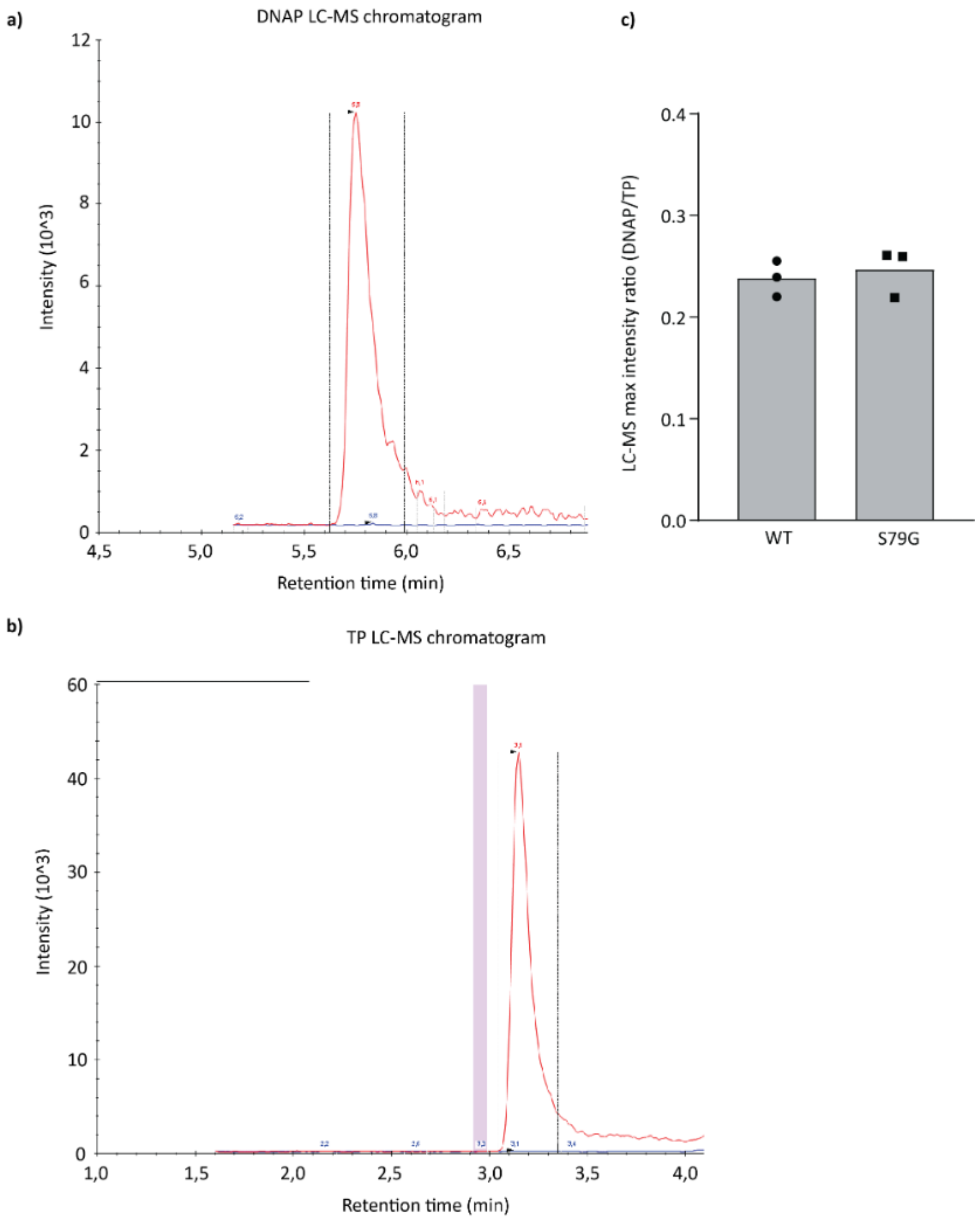


**Fig. S14: Wild-type and S79G DNAP proteins are expressed at similar levels in PURE system. a,b)** LC-MS chromatograms of pre-ran PURE samples containing expressed DNAP and TP. **c)** Relative quantification of DNAP and TP was obtained by normalizing the peak intensity for DNAP (a), either WT or S79G, with the peak intensity of TP (b). Each symbol represents a separate experiment and the bar height is the average of the three repeats.

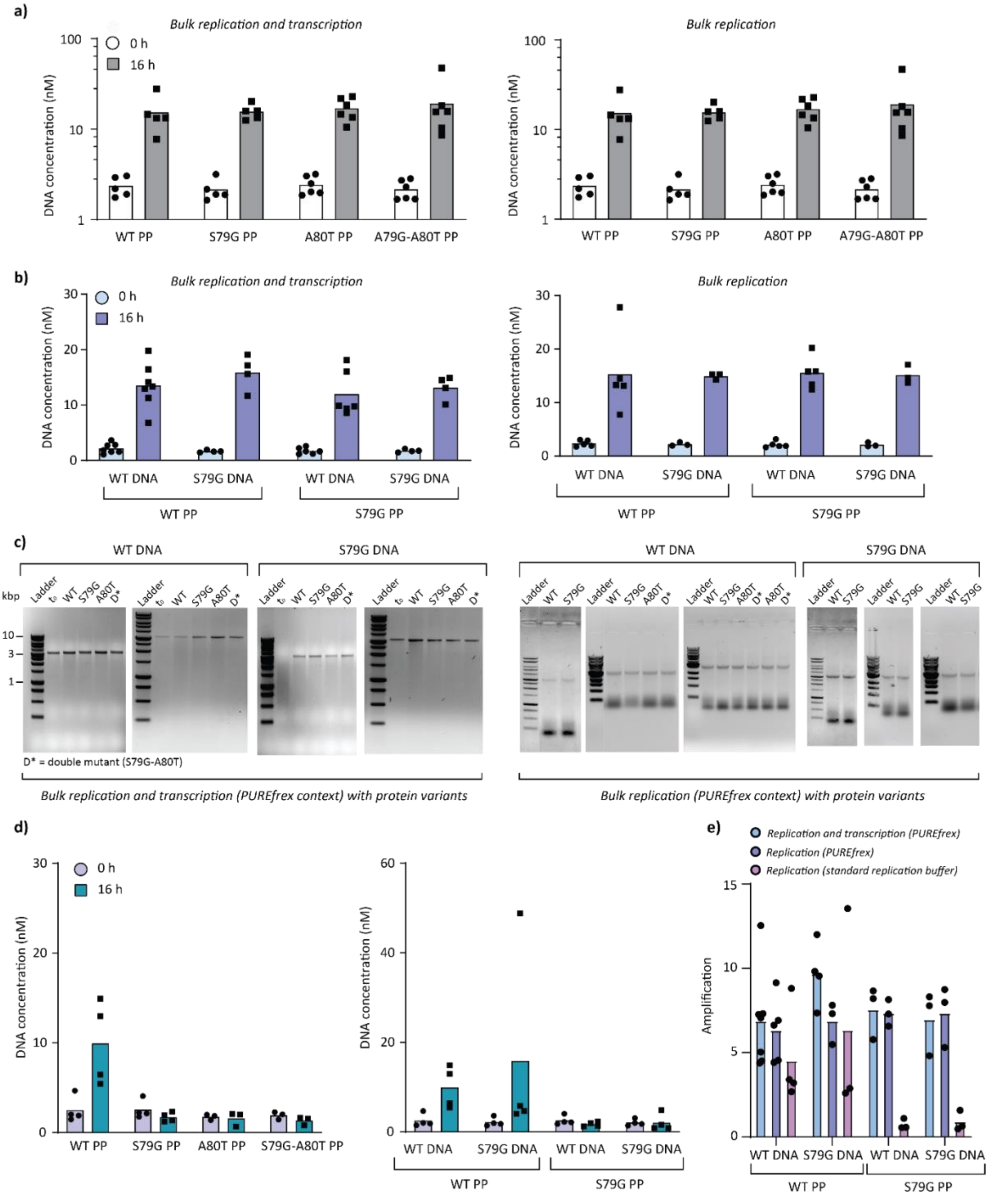


**Fig. S15: Bulk replication assays of *mod-ori-p2p3* templates with purified DNAP protein variants: S79G, A80T, and double DNAP mutant. a)** DNA concentration in bulk replication-transcription (left) and replication (right) samples with different DNAP variants, as indicated, in a PURE*frex* background. **b)** DNA concentration in bulk replication-transcription (left) or replication (right) samples with different DNA templates: *mod-ori-p2p3* or *mod-ori-p2(S79G)p3*, and different purified DNAP in a PURE*frex* background. Similar replication efficiency is observed for all tested conditions**. c)** Agarose gel electrophoresis of *mod-ori-p2p3* and *mod-ori-p2(S79G)p3* recovered DNA from the reaction samples shown in panels (a) and (b). The DNA template is indicated on the square brackets and the protein variants on the gel lanes. **d)** Bulk replication reactions were performed in a standard Φ29 DNA replication buffer using either DNA template *mod-ori-p2p3* or *mod-ori-p2(S79G)p3*, and different purified DNAP variants, as indicated. In the left panel, *mod-ori-p2p3* was used. **e)** Amplification (DNA concentration at 16 h /

DNA concentration at 0 h) for the different conditions tested. Individual data points per condition correspond to biological repeats. DNA concentration was determined by qPCR. PP, purified protein.

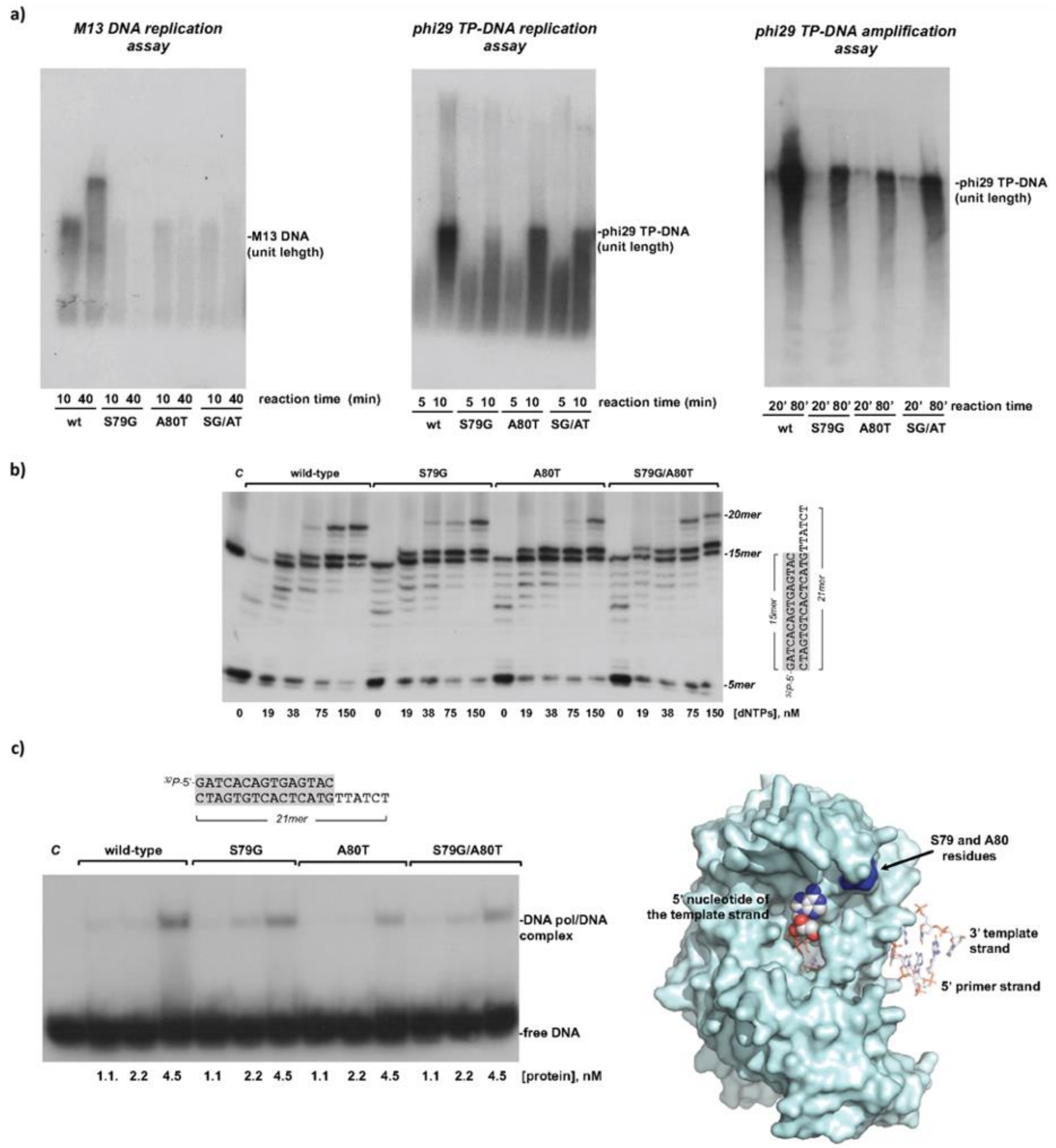


**Fig. S16: Replication assays with purified DNAP variants. a)** M13 DNA replication assay (left). Replication of primed M13 DNA was carried out as described in Materials and Methods in the presence of 60 nM of either wild-type or mutant DNA polymerase. The position of full-length M13 DNA is shown at the right of the gel. TP-DNA replication assay (middle). The assay was performed essentially as described in Materials and Methods using 12 nM of TP and 12 nM of either wild-type or mutant DNA polymerase. The migration position of unit-length of TP-DNA is indicated. DNA amplification assay (right). The assay was carried out as described in Material and Methods, in the presence of 3 nM of DNA polymerase wild-type or the indicated mutant, 6 nM of TP, 30 µM of SSB and 30 µM of DBP. The migration position of unit-length of TP-DNA is indicated. SG/AT denotes the double mutant S79G/A80T. **b)** Exonuclease/polymerisation balance assay. The reaction was performed essentially as described in Materials and Methods using as substrate the 5' labelled molecule sp1/sp1c+6 (depicted at the right of the figure), 30 nM of either wild-type or mutant DNA polymerase and the indicated concentration of nucleotides. Polymerisation or 3'-5' exonuclease activity is detected as an increase or decrease in the size of the labelled primer (15 mer). *C*: control lane without enzyme. **c)** Interaction of wild-type and mutant DNA polymerases with a primer/template substrate. The EMSA assay was performed as described in Materials and Methods. The 5' labelled molecule sp1/sp1c+6, 15mer/21mer (depicted at the top of the figure) was incubated either with the wild-type or mutant enzymes. The bands corresponding to free DNA or to the DNA/DNA polymerase complex were detected by autoradiography. *C*: control lane without enzyme. Surface representation of Φ29 DNA polymerase complexed with primer/template DNA (right). Crystallographic data are from Protein Data Bank ID code 2PZS. DNA polymerase residues S79 and A80 are shown as spheres and the primer and template strands as sticks. Figure was generated using The Open-Source Molecular Graphics System, v. 2.5.0, Schrödinger, LLC (Open-Source PyMOL is Copyright (C) Schrodinger, LLC.).
